# Modulation by neuropeptides with overlapping targets results in functional overlap in oscillatory circuit activation

**DOI:** 10.1101/2023.06.05.543756

**Authors:** Elizabeth M. Cronin, Anna C. Schneider, Farzan Nadim, Dirk Bucher

## Abstract

Neuromodulation lends flexibility to neural circuit operation but the general notion that different neuromodulators sculpt neural circuit activity into distinct and characteristic patterns is complicated by interindividual variability. In addition, some neuromodulators converge onto the same signaling pathways, with similar effects on neurons and synapses. We compared the effects of three neuropeptides on the rhythmic pyloric circuit in the crab *Cancer borealis* stomatogastric nervous system. Proctolin (PROC), crustacean cardioactive peptide (CCAP), and red pigment concentrating hormone (RPCH) all activate the same modulatory inward current, *I*_MI_, and have convergent actions on synapses. However, while PROC targets all four neuron types in the core pyloric circuit, CCAP and RPCH target the same subset of only two neurons. After removal of spontaneous neuromodulator release, none of the neuropeptides restored the control cycle frequency, but all restored the relative timing between neuron types. Consequently, differences between neuropeptide effects were mainly found in the spiking activity of different neuron types. We performed statistical comparisons using the Euclidean distance in the multidimensional space of normalized output attributes to obtain a single measure of difference between modulatory states. Across preparations, circuit output in PROC was distinguishable from CCAP and RPCH, but CCAP and RPCH were not distinguishable from each other. However, we argue that even between PROC and the other two neuropeptides, population data overlapped enough to prevent reliable identification of individual output patterns as characteristic for a specific neuropeptide. We confirmed this notion by showing that blind classifications by machine learning algorithms were only moderately successful.

**Significance Statement:** It is commonly assumed that distinct behaviors or circuit activities can be elicited by different neuromodulators. Yet it is unknown to what extent these characteristic actions remain distinct across individuals. We use a well-studied circuit model of neuromodulation to examine the effects of three neuropeptides, each known to produce a distinct activity pattern in controlled studies. We find that, when compared across individuals, the three peptides elicit activity patterns that are either statistically indistinguishable or show too much overlap to be labeled characteristic. We ascribe this to interindividual variability and overlapping subcellular actions of the modulators. Because both factors are common in all neural circuits, these findings have broad significance for understanding the chemical neuromodulatory actions while considering interindividual variability.

## Introduction

The flexibility of neural circuit operation that allows adaptation to different behavioral contexts is largely owed to neuromodulation (Briggman and Kristan, 2008; Doi and Ramirez, 2008; Bargmann, 2012; Marder, 2012; Bargmann and Marder, 2013; Bucher and Marder, 2013; Nadim and Bucher, 2014; Diaz-Rios et al., 2017; McCormick et al., 2020). Consequently, specific types of circuit activity, and entire brain states, have been associated with specific neuromodulators. The assumption is that different neuromodulators, alone or in combination, elicit characteristic and distinguishable activity. This is straightforward to validate when different neuromodulators are associated with qualitatively different behaviors. For example, crawling and swimming in leech and *C. elegans* are associated with dopamine and serotonin, respectively (Briggman and Kristan, 2006; Vidal-Gadea et al., 2011), and sleep-wake cycles in mammals are controlled by multiple neuromodulators (Fuller et al., 2006; Jones, 2020). It is less straightforward when neuromodulators push circuits across a continuous spectrum of activity, without clear transitions between distinct behaviors. For example, circuits controlling locomotion can execute the same gait at different speeds (Grillner, 2006; Mullins et al., 2011; Zhang et al., 2014; Le Gal et al., 2017). In addition, the question of what constitutes circuit activity characteristic for a specific modulator is complicated by convergent actions of different neuromodulators, and by interindividual variability.

Neuromodulators, principally amines and neuropeptides, act mostly through G-protein-coupled receptors and alter excitability and synaptic function in a cell type-specific manner. Because their release is often paracrine or hormonal, they affect multiple neurons at once and sculpt circuit activity dependent on patterns of convergence and divergence in their cellular actions (McCormick and Williamson, 1989; Harris-Warrick and Johnson, 2010; Harris-Warrick, 2011; Bucher and Marder, 2013; Nadim and Bucher, 2014; Due et al., 2022). Two neuromodulators can target different but overlapping subsets of neurons. In a single neuron responsive to both, effects on ion channels or synaptic release can be divergent or convergent. If receptor activation converges on the same signaling pathway in enough neurons, circuit effects may be similar and only differ in the context of release (McCormick and Williamson, 1989). In addition, even divergent cellular mechanisms can give rise to qualitatively similar circuit activity (Saideman et al., 2007; Rodriguez et al., 2013).

Another confounding factor for consistent neuromodulator effects is interindividual variability. Ionic currents and synapses targeted by neuromodulators can vary substantially in magnitude and strength, and circuit outputs under the same conditions can show some degree of variability (Marder and Goaillard, 2006; Marder et al., 2007; Calabrese et al., 2011; Golowasch, 2014; Marder et al., 2014a; Goaillard and Marder, 2021). Modulator release and receptor expression can also be quite variable (Garcia et al., 2015; Maloney, 2021). Interestingly, through a range of different mechanisms, neuromodulation can either promote or counteract interindividual variability in neuron and circuit activity (Hamood and Marder, 2014; Marder et al., 2014b; Maloney, 2021; Xu et al., 2021; Schneider et al., 2022; Tamvacakis et al., 2022; Schneider et al., 2023).

Here we ask to which degree different neuromodulators with convergent cellular actions, but divergent circuit targets produce reliably distinguishable activity across individuals. We compare activity of the pyloric central pattern generator of the crustacean stomatogastric ganglion (Marder and Bucher, 2007; Stein, 2009; Daur et al., 2016) across bath applications of different neuropeptides. While in this system modulatory effects are tractable from cellular to circuit level, few studies have directly compared actions of different neuromodulators on circuit output (Marder and Weimann, 1992). The principal action of these neuropeptides is the activation of a voltage-gated inward current, *I*_MI_ (Golowasch and Marder, 1992; Swensen and Marder, 2000; Gray and Golowasch, 2016; Gray et al., 2017). Despite this convergence, they are thought to produce distinct circuit outputs because each targets an overlapping but different subset of neurons (Swensen and Marder, 2001). However, we show that across a population of individuals with varying responses, even statistically distinguishable output is not sufficiently characteristic to allow identification of neuromodulatory state from individual recordings.

## Materials and Methods

### Experimental preparation

All experiments were performed on adult male crabs (*Cancer borealis*) obtained from local fish markets and maintained in recirculating artificial seawater tanks at 12°C until use. Crabs were anesthetized in ice for ∼30 minutes prior to dissection. Dissections were performed in saline containing: 11 mM KCl, 440 mM NaCl, 13 mM CaCl_2_ · 2H_2_O, 26 mM MgCl_2_ · 6H_2_O, 11.2 mM Tris base, 5.1 mM maleic acid, with a pH of 7.4. The stomatogastric nervous system (STNS), including the stomatogastric ganglion (STG), esophageal ganglion (OG), the pair of commissural ganglia (CoG), and the motor nerves (schematic in Figure 1A) were dissected from the stomach and pinned to a saline-filled, Sylgard-coated (Dow Corning) Petri dish. The STG was desheathed to facilitate intracellular impalement of neuronal somata and allow for easy permeation of bath applied neuromodulators. Preparations were superfused with chilled (11-13°C) physiological saline using a peristaltic pump (Dynamax).

**Figure 1:**
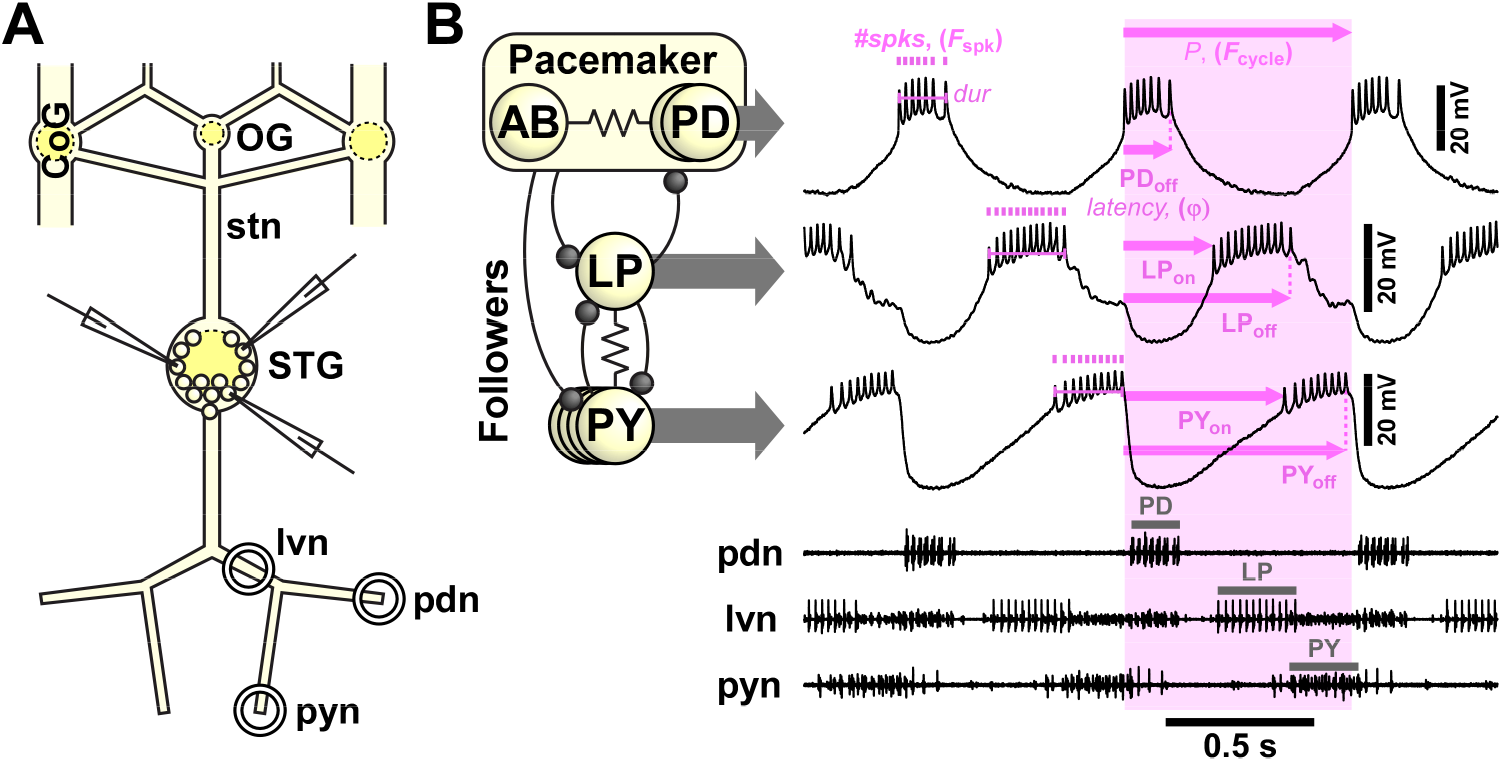
Experimental preparation and example recording traces. **A:** Schematic of the isolated STNS with three intracellular recording electrodes in the STG. CoG: commissural ganglion; OG: esophageal ganglion; stn: stomatogastric nerve; STG: stomatogastric ganglion; lvn: lateral ventricular nerve; pdn: pyloric dilator nerve; pyn: pyloric nerve. Nomenclature after Maynard and Dando (1974). **B:** Simplified pyloric circuit diagram (inset) and example traces from simultaneous intracellular soma recordings and extracellular nerve recordings, indicating the output pattern attributes used for analyses. Ball-and-stick connections in inset indicate synaptic inhibition, whereas resistor symbols indicate electrical coupling. Multiple circles indicate multiple copies of that neuron type.

### Electrophysiological recordings

In this study we focus on the core subcircuit of the pyloric circuit which consists of 4 neuron types that have a well-characterized connectivity pattern (Fig. 1B). The two pyloric dilator (PD) neurons are motor neurons that are electrically coupled to each other and to the anterior burster (AB) neuron. The PD and AB neurons form the pacemaker kernel of the pyloric central pattern generator. The follower neurons, including the single lateral pyloric (LP) neuron and the 3-5 pyloric (PY) constrictor neurons, sequentially burst in rebound from inhibition by the pacemakers. The pyloric rhythm is therefore a triphasic bursting pattern (Marder and Bucher, 2007) (Fig. 1B). The activity of the PD, LP, and PY neurons was recorded simultaneously with multiple intracellular soma recordings and extracellular motor nerve recordings.

Extracellular recordings from identified motor nerves were obtained with pairs of stainless-steel wire electrodes, placed inside and outside of petroleum jelly wells built to electrically isolate small sections of nerve. Extracellular signals were amplified using differential AC amplifiers (A-M Systems, model 1700). PD activity was recorded from the pyloric dilator nerve (pdn). LP activity was recorded from the lateral ventricular nerve (lvn), where LP contributes the signal with the largest amplitude. PY activity was recorded from the pyloric constrictor nerve (pyn). Intracellular recordings were obtained from neuronal somata in the STG using sharp glass microelectrodes prepared with the Flaming-Brown micropipette puller (P97, Sutter Instruments). They were filled with an electrolyte solution of 0.6 M K_2_SO_4_ and 20 mM KCl, yielding tip resistances of 20-30 MΩ. Electrodes were positioned with MicroStar motorized manipulators (Scientifica). Intracellular signals were amplified with Axoclamp 2B and 900A amplifiers (Molecular Devices). Neurons were identified by their characteristic waveforms and by matching their spike patterns to the extracellular activities recorded in the corresponding motor nerves. Both intracellular and extracellular signals were digitized at 10 kHz using a Micro1401 digitizer with its accompanying Spike2 data acquisition software (Cambridge Electronic Design).

### Neuromodulator application

In all preparations, petroleum jelly wells were built around the STG for focal neuromodulator superfusion. In addition, a well was built around the stomatogastric nerve (stn), the sole input nerve to the STG from other central nervous system ganglia. The triphasic pyloric rhythm was recorded first with intact descending modulatory inputs from the CoGs and OG to the STG. Subsequently, the STG was decentralized, i.e., modulatory inputs were removed by blocking action potential conduction in descending axons. This was achieved by application of 100 nM TTX (tetrodotoxin citrate, Biotium) in a 750 mM sucrose (Sigma) solution to the well around the stn. Neuromodulators were then applied to the STG at a concentration of 1 μM for at least 5 min. When multiple modulators were applied sequentially to the same preparation, each modulator was washed out for at least 30 min. These modulators included DA (dopamine: 3-hydroxytyramine hydrochloride, Sigma) and the neuropeptides PROC (proctolin, custom synthesis by RS synthesis), CCAP (crustacean cardioactive peptide, Bachem), and RPCH (red pigment concentrating hormone, Bachem). DA stock solutions of 1 mM in distilled water were always made fresh and kept in the dark on ice for no longer than 1 h. PROC and CCAP were initially dissolved in distilled water, and RPCH in dimethyl sulfoxide (DMSO, Sigma), to yield stock solutions of 1 mM, which were then aliquoted and held at -20°C until needed. Final dilution of modulators to the experimental concentration occurred immediately before application in chilled saline.

### Initial pattern analysis

Initial analysis of circuit activity patterns was performed using custom-written scripts in Spike2. Cycle-to-cycle spike and burst detection allowed extraction of several output pattern attributes (Fig. 1B). Cycle period (*P*) was defined as the latency between consecutive PD burst starts. Cycle frequency (*F*_cycle_) was then calculated as 1/*P*. Using the PD burst start as the reference time, we measured the latencies of all burst starts (on) and burst ends (off), and burst durations as the time between burst starts and burst ends. Latencies were then divided by *P* to yield values for phase (ϕ). The numbers of spikes per burst (*#spks*) were determined for all neurons, and the spike frequencies within bursts (*F*_spk_) were calculated as *#spks*-1 divided by the burst duration. From each individual recording and state, we used a minimum of 20 cycles and determined the mean value for each attribute.

While all output attributes are potentially interdependent, some are unambiguously redundant. Therefore, we only used a subset of attributes for our analyses. *P* and *F*_cycle_ are the inverse of each other, so we only used *F*_cycle_. As latencies and burst durations were strongly dependent on *P*, we only used the relative timing (ϕ_on_ and ϕ_off_). We also excluded duty cycle (burst duration/*P*), as it is merely the difference between ϕ_off_ and ϕ_on_. *F*_spk_ and *#spks* are obviously interdependent. However, they were not strongly correlated in at least a subset of neurons and conditions (e.g., LP and PY in neuropeptides showing R^2^ values < 0.5). Therefore, we used both.

In extracellular recordings, spikes from the two PD axons in the pdn and from multiple PY axons in the pyn summate, which renders individual spike detection ambiguous. Therefore, all analyses that included the full set of rhythm attributes were performed only from experiments in which we established simultaneous intracellular recordings from PD, LP and PY neurons. Extracellular recordings were only used for additional analyses of burst timing, as indicated in the Results section. Basic descriptive statistics of pattern attributes and comparisons across modulator treatments with analysis of variance (ANOVA) and subsequent Holm-Sidak *post-hoc* tests were performed using SigmaPlot (version 12.0, Systat Software). All subsequent data handling and analysis described in the following was performed in MATLAB (version 2022a, Mathworks). Statistical significance was assumed at *p* < 0.05 and is indicated as * *p* < 0.05, ** *p* < 0.01, and *** *p* < 0.001.

### Euclidian distance and multivariate permutation tests

For comparisons between modulatory conditions in multiple dimensions, we determined the mean of each attribute across experiments and calculated the Euclidian distance:

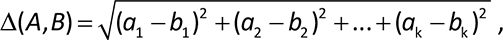

where *A* and *B* are two modulatory conditions, and *a*_1_ to *a*_k_ and *b*_1_ to *b*_k_ are the means of the different output pattern attributes as Cartesian coordinates in k-dimensional Euclidian space.

We did this first with attributes of the same units across different neurons: Δϕ as the distance in 5 dimensions (ϕPD_off_, ϕLP_on_, ϕLP_off_, ϕPY_on_, ϕPY_off_), Δ*#spks* in 3 dimensions (PD *#spks*, LP *#spks*, PY *#spks*), and Δ*F*_spk_ in 3 dimensions (PD *F*_spk_, LP *F*_spk_, PY *F*_spk_). To be able to include attributes of different units in the same analysis, we standardized each attribute by calculating *z*-scores:

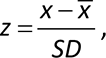

where *x* is an attribute in a single experiment, and 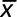 and *SD* are, respectively, the mean and standard deviation of that attribute across all experiments and modulatory states.

Once all attributes were z-scored, we calculated the Euclidian distance between pairs of conditions in 12 dimensions (*F*_cycle_, 5 dimensions of ϕ, 3 dimensions of *#spks*, 3 dimensions of *F*_spk_). Principal component analysis (PCA) was used to visualize the 12-dimensional data in the three-dimensional space of the first three principal components.

To determine if differences between two modulatory states were statistically significant, we used a modified version of the multi-variate permutation test (Anderson, 2001; Nichols and Holmes, 2002). For example, to compare data from modulatory states *A* and *B* with respective sample sizes n*A* and n*B* (each sample with k coordinates), we first calculated the Euclidean distance between the means of the two modulatory states, as described above. The data from the two groups were then pooled into a single group, and data points were randomly reassigned into two new groups, *A*i and *B*i, of the same respective sizes (n*A* and n*B*) by exchanging (permuting) elements between the two groups. We then calculated the Euclidean distance between the means of the two new groups *A*i and *B*i. This process was repeated for 9999 permutations of *A*i and *B*i, and the one-sided p-value of the test was calculated as the proportion of permutations for which the Euclidean distances were larger than the original distance of A and B. The determination of statistical significance was made with an alpha level of 0.05.

### Machine learning analysis

As an independent test for how well output patterns from different modulatory conditions can be distinguished, we used supervised machine learning (ML) algorithms. Pattern recognition and time series classification networks were created using the Deep Learning Toolbox (version 14.2) and the Parallel Computing Toolbox (version 7.4) in MATLAB.

Because training outcome partially depends on initial values of ML network parameters and our datasets were relatively small for this purpose, we used a bootstrapping approach to sample the distribution of classification accuracy based on network initialization and dataset splitting in training and test data. For each modulatory condition, we split the corresponding dataset randomly into 70% training data and 30% test data. We then randomly initialized 50 ML networks, trained them, and kept the most accurate one. On the next run, we again split the dataset randomly into training and test data and repeated the same process. We repeated this process for 100 runs, which resulted in 100 networks of varying accuracy. We then classified the test data with the corresponding networks and calculated the percentage of correct and false positive classifications across these 100 ML networks for each modulatory condition.

We first tested this approach with the *z*-scored attributes that we had extracted for our Euclidian distance analysis. Because all temporal features of output activity are explicit (already quantified) in these attributes, we could employ a shallow feedforward pattern recognition network (Janiesch et al., 2021). This ML algorithm was created with the default settings of the MATLAB function “patternnet” and consisted of one input layer, one hidden layer with 50 units, and one output layer.

We also performed a different ML analysis based only on (not *z*-scored) spike times and *F*_cycle_. For this, we extracted spike times of the three neurons (PD, LP, and PY) from 30 s of recording data from each trial in each experiment and sorted them into a fixed number of bins per cycle, thereby generating spike phase histograms. To account for differences in *F*_cycle_ across modulatory states, we optionally biased the bin counts by adding the *z*-scored values of *F*_cycle_. The more minimally processed data does not explicitly contain relative timing information and therefore requires an ML network that is able to process temporal sequences within and across neurons. Therefore, we resorted to using a time series classification network (Goodfellow et al., 2016). This recurrent neural network consisted of a sequence input layer with three channels (one for each neuron), a bi-directional long short-term memory layer (LSTM, Schuster and Paliwal (1997)) with 50 hidden units set to output the last time step of the sequence, a fully connected layer with five channels (one for each modulatory state), a softmax layer to convert real numbers into probability distributions, and the classification layer. We empirically determined that in order to obtain trained networks with satisfactory accuracy, we had to change the default settings in MATLAB to use the Adam optimizer with a denominator offset of 10^-6^, set the mini-batch size to accommodate all input data in one batch, allow a maximum of 1000 epochs for training, validate the network at every other iteration with the test data, and set the patience of validation stopping to 25. It would have been possible to use the original voltage traces for time series classification with this network (Janiesch et al., 2021), but the minimal processing reduced the amount of data sufficiently to train the network on a desktop PC in a reasonable amount of time.

To visualize the classification results from the ML networks, we show classification matrices modified from the commonly used confusion matrices to account for the unequal numbers of recordings from each modulatory state. The matrices show the actual modulatory state as the true class for each row, and the percentage of correct and incorrect classifications in each column for that row. Therefore, percentages add up to 100% in each row instead of for the whole matrix.

## Results

### Circuit outputs in different neuropeptides were distinguishable by spiking behavior but not burst timing

We characterized the pyloric rhythm by determining output pattern attributes from simultaneous recordings of PD, LP, and PY neurons (Fig. 1B). Intracellular recordings showed slow wave oscillations and attenuated spikes typical for soma recordings of STG neurons, with the well-characterized triphasic sequence between neurons (Marder and Bucher, 2007). We measured attributes across different modulatory states, with intact descending input, after decentralization, and after application of neuropeptides (Fig. 2A). We assumed that with intact descending modulatory inputs, all neurons and synapses were modulated, while decentralization removed all modulatory control of intrinsic and synaptic properties. PROC application activates *I*_MI_ in AB, PD, LP, and ∼70% of PY neurons, while both CCAP and RPCH activate *I*_MI_ only in AB and LP (Swensen and Marder, 2001). Both PROC and CCAP enhance synaptic currents in the reciprocal connections between the pacemaker neurons and LP (Li et al., 2018), and RPCH enhances the LP to pacemaker synaptic current (Atamturktur and Nadim, 2011), but effects on the pacemaker to PY and PY to LP synapses have not been studied.

**Figure 2:**
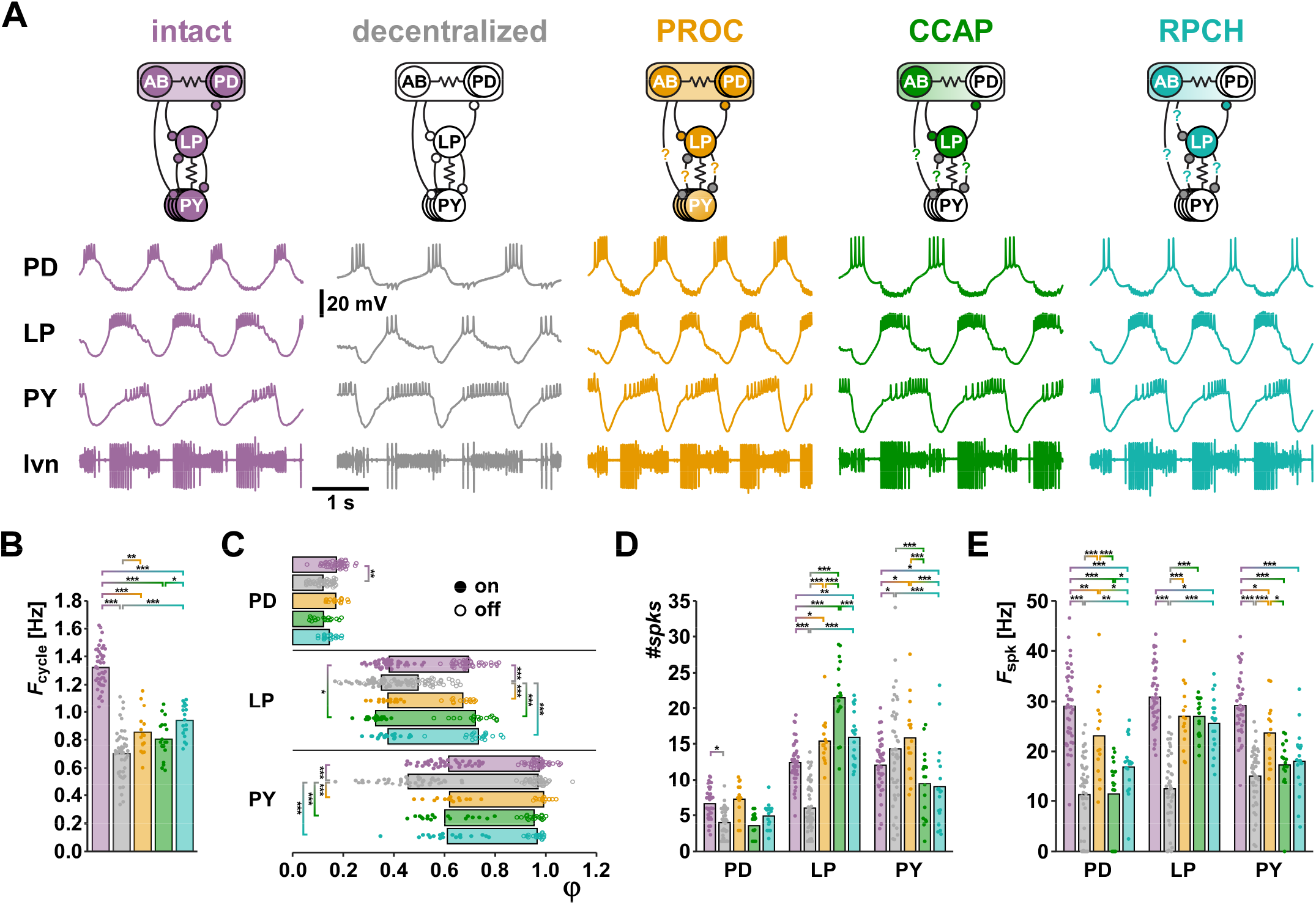
Comparison of circuit output patterns under different neuromodulatory conditions. **A:** Example traces from a single preparation under different modulatory conditions. In the circuit diagrams, neurons and synapses are shown in color if they are modulated in a given condition, and in white if they are not modulated (see main text for references). Synaptic connections for which there is no available information about modulation are gray and marked with question marks. **B:** Means (bars) and individual data points (circles) for *F*_cycle_ (intact: n = 43; decentralized: n = 46; PROC: n = 15; CCAP: n = 18; RPCH: n = 19). One-way ANOVA showed significant difference between treatments (F_(4,140)_ = 101.3, p < 0.001, power = 1, normality and equal variance tests passed). Asterisks indicate significant difference determined by subsequent Holm-Sidak post-hoc tests. **C:** Mean values of ϕ (bars) and individual data points (filled circles: ϕ_on_, open circles: ϕ_off_). Two-way ANOVA showed significant interactions (F_(16,704)_ = 15.7, p < 0.001, power = 1, normality and equal variance tests failed). Asterisks indicate significant difference determined by subsequent Holm-Sidak post-hoc tests. **D:** Means (bars) and individual data points (circles) for *#spks*. Two-way ANOVA showed significant interactions (F_(8,422)_ = 26.1, p < 0.001, power = 1, normality and equal variance tests failed). Asterisks indicate significant difference determined by subsequent Holm-Sidak post-hoc tests. **E:** Means (bars) and individual data points (circles) for *F*_spk_. Two-way ANOVA showed significant interactions (F_(8,422)_ = 6.4, p < 0.001, power = 1, normality and equal variance tests passed). Asterisks indicate significant difference determined by subsequent Holm-Sidak post-hoc tests.

We first used the measured output pattern attributes for standard statistical evaluation (Fig. 2B-E). In most cases, the required long washes between neuropeptide applications prevented us from obtaining data from all modulatory states in the same experiment. Therefore, all statistics were performed as unpaired comparisons. Decentralization substantially slowed the pyloric rhythm and none of the neuropeptides on their own recovered *F*_cycle_ to the intact value (Fig. 2B). In fact, only in RPCH was *F*_cycle_ significantly increased from the decentralized state. *F*_cycle_ in RPCH was also larger than in CCAP, but not distinguishable from PROC. The activity phases of the three neurons were changed substantially by decentralization, but all neuropeptides essentially restored the phase relationships to the intact state (Fig. 2C). Only ϕLP_on_ in CCAP was different from the intact pattern. In contrast, the three neuropeptides had several differential effects on *#spks* and *F*_spk_ (Fig. 2D, E). LP *#spks* differed between PROC and CCAP and between CCAP and RPCH. PY *#spks* in PROC differed from both CCAP and RPCH. PD *F*_spk_ differed between all peptides, and PY *F*_spk_ between PROC and CCAP. It therefore appears that, at least at first pass, the modulated states should be more easily distinguishable based on spiking activity within bursts than based on *F*_cycle_ or phase relationships.

### Covariance showed interdependence of circuit output attributes

The type of analysis shown in Figure 2 comes with some caveats. In recurrent circuits, all activity attributes within and across different neuron types potentially depend on each other. To establish an effect of a neuromodulator, it is appropriate to ask whether the modulator changes a given single attribute like *F*_cycle_ or *#spks* in a specific neuron. However, when many attributes are separately tested across two or more circuit states, and in several neurons, the interdependence of these attributes is at least partially ignored. Statistical tests like ANOVA may report the presence of such interdependence but can only be used for attributes within the same category (e.g., all activity phases).

Such interdependence is illustrated by the relationships between *P* (or equivalently, *F*_cycle_) and burst timing. In the intact pyloric circuit, latencies of burst on and off times scale with *P*, so that values of ϕ are largely independent of *F*_cycle_. This is true both within individuals when *F*_cycle_ is experimentally altered (Hooper, 1997; Tang et al., 2012; Soofi et al., 2014), and across individuals that differ in *F*_cycle_ under control conditions (Bucher et al., 2005; Goaillard et al., 2009; Anwar et al., 2022). We found that phases are also largely independent of frequencies across different modulated states. In pooled data from all modulated conditions (intact, PROC, CCAP, RPCH), latencies scaled roughly proportionally with *P* across all states (Fig. 3A, top left panel). Linear regressions showed small offsets and large R^2^ values (Fig. 3-1). Consequently, values of ϕ were mostly independent of *F*_cycle_ (Fig. 3A, top right panel). In contrast, latencies did not scale proportionally with *P* in decentralized preparations, rendering ϕ values dependent on *F*_cycle_. (Fig. 3A, bottom panels) The only exception was PY_off_, as PY neurons usually burst until the end of each cycle when they are inhibited by the next pacemaker burst (i.e., ϕPY_off_ is always close to 1).

**Figure 3:**
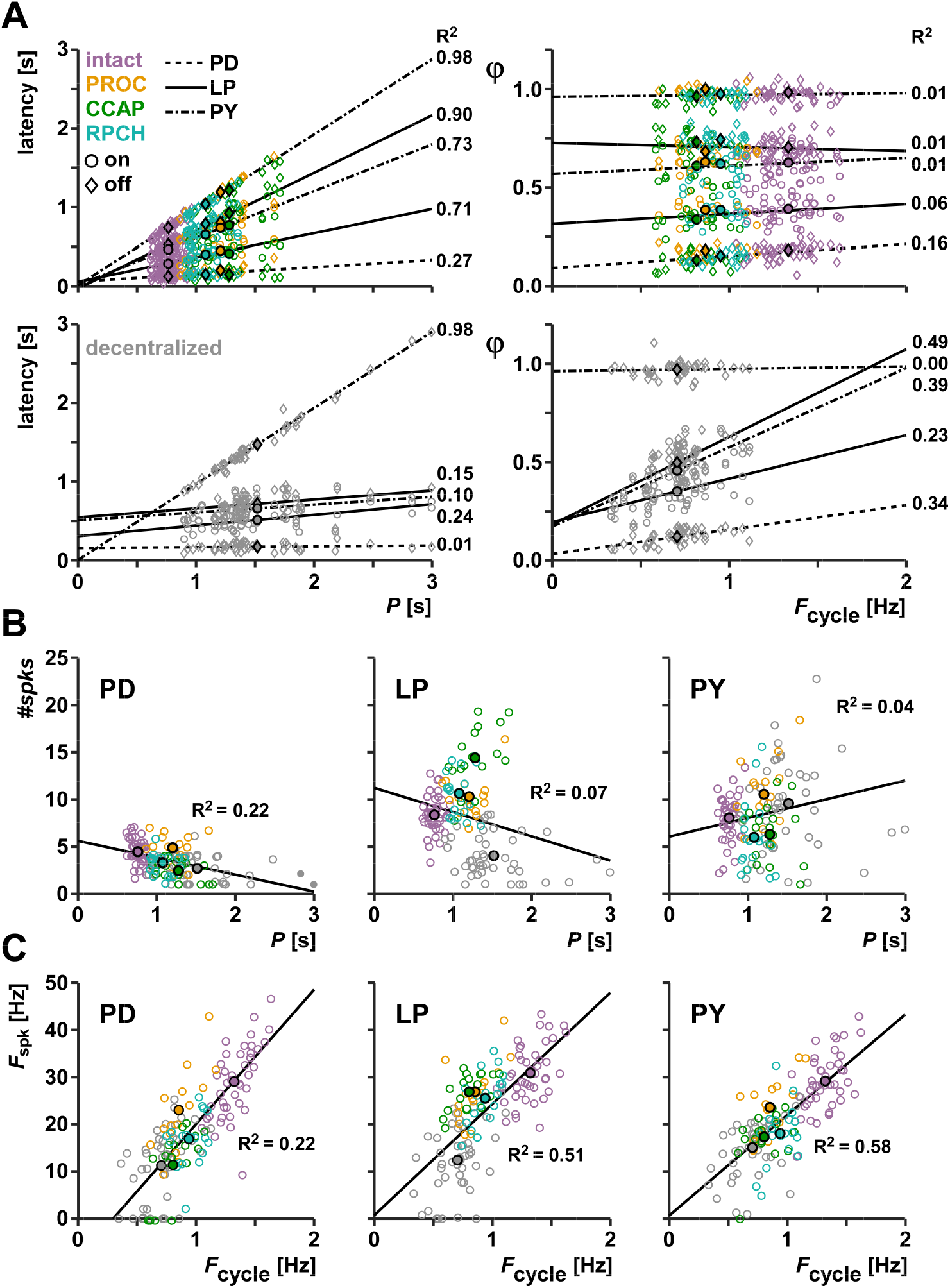
Dependence of burst and spike attributes on *P* or *F*_cycle_. **A:** Plot and regression of latencies vs. *P*, and ϕ vs. *F*_cycle_, for all the combined modulated states (intact, PROC, CCAP, RPCH; upper panels) and the decentralized state (lower panels). Points with black edges indicate mean values. **B:** Regression of *#spks* vs. *P* for all combined states in the three neuron types. **C:** Regression of *F*_spk_ vs. *F*_cycle_ for all combined states in the three neuron types. **Figure 3-1**: Regression parameters.

We also analyzed the relationship between *P* or *F*_cycle_ and spiking activity (Fig. 3B, C and Fig. 3-1). Across all states, values of *#spks* change with *P* in all neurons (Fig. 3B). However, this scaling was not proportional, and coefficients of determination were small. *F*_spk_ across all states was dependent on *F*_cycle_ in all neurons (Fig. 3C), and *F*_spk_ in LP and PY scaled proportionately with *F*_cycle_. In this case, R^2^ values were larger than for the relationship between *#spks* and *P*, but smaller than for the relationship of latencies and *P*.

These results suggest that the constant phases across *F*_cycle_ arise from a stable dependence of latency on *P* that is independent of which neuropeptide is present, while spiking behavior within bursts is more malleable.

### Multidimensional comparison confirmed the similarity of phase relationships in different modulated conditions

In addition to interdependence of attributes, quantification of differences is usually only established one-by-one for each attribute. For our analysis, however, we wanted to ask how different or similar the entire output patterns are across different modulatory states, preferably with a method that yields a single measure of difference between each pair of modulatory states. To this end, we measured the distance between circuit outputs in the multidimensional attribute space. As proof-of-concept, we first did this type of analysis separately for attributes with the same units.

We first examined the effects of the modulators in the five-dimensional ϕ space. For ease of visualization, we plotted two different combinations, each showing three of the five ϕ values (LP_on_, LP_off_, PY_on_ in Fig. 4A; PD_off_, LP_off_, PY_off_ in Fig. 4B). The distribution of data points from each individual experiment (Fig. 4Ai, Bi) showed substantial overlap between the modulated states (intact, PROC, CCAP, RPCH) but much smaller overlap between the decentralized state and all modulated states. To visualize the mean differences (Fig. 4Aii, Bii), we plotted the mean Cartesian coordinates (centroid) for each state as a sphere and connected each pair of centroids with a line representing the three-dimensional Euclidian distance. In both combinations of phases, the centroids of the modulated states clustered while the centroid of the decentralized state was more distant, resulting from differences in all dimensions but PY_off_.

**Figure 4:**
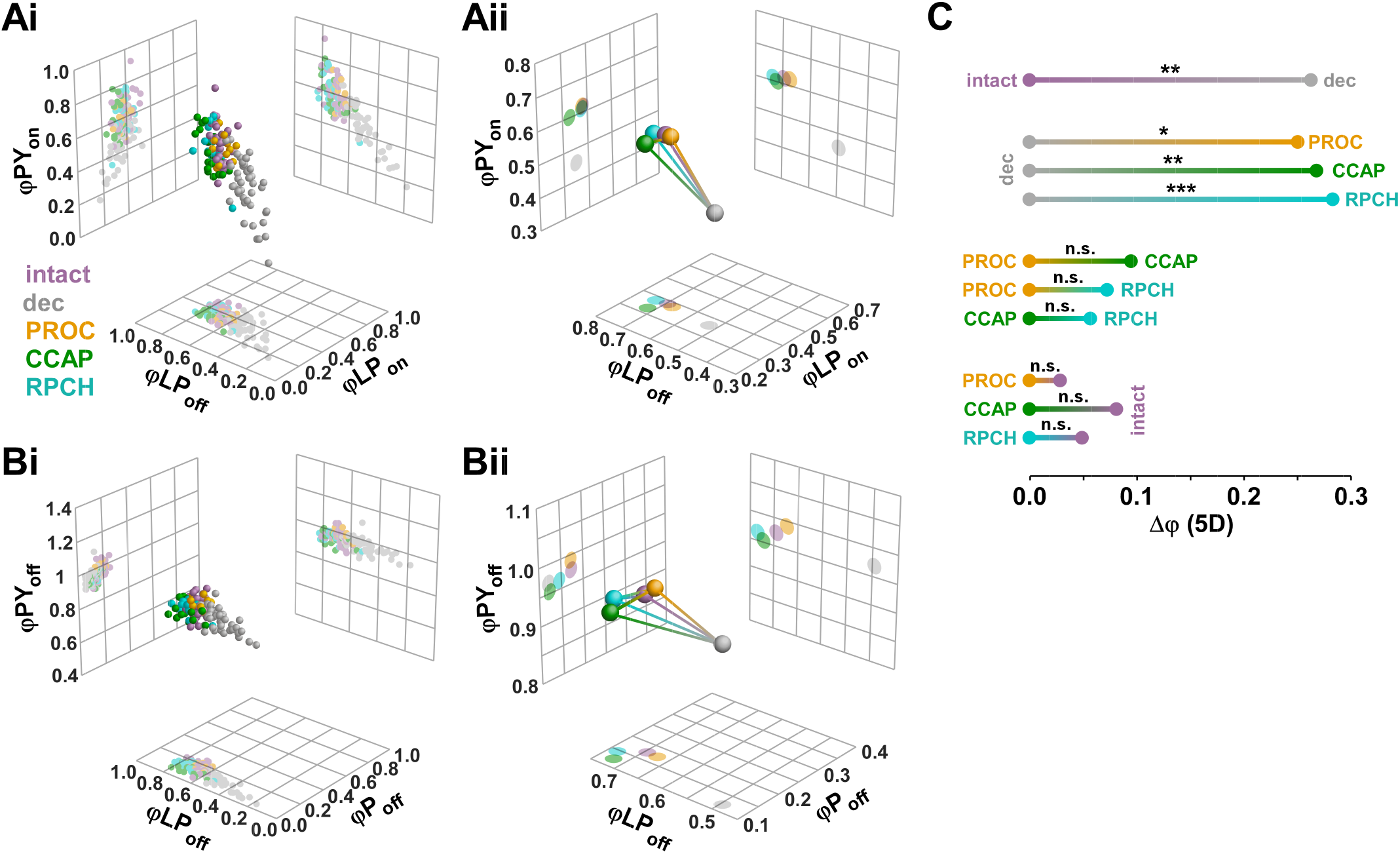
Euclidian distance analysis of phase relationships. Data set is the same as shown in Figure 2. **A:** Three-dimensional plots (spheres) and corresponding two-dimensional projections (grids) of three of the five phases (ϕLP_on_, ϕLP_off_, and ϕPY_on_), for individual experiments (i) and centroids (ii). Lines connecting the centroids represent the Euclidian distances in three dimensions. **B:** Plots of a different combinations of three of the five phases (ϕPD_off_, ϕLP_off_, ϕPY_off_). **C:** One-dimensional projections of the five-dimensional Euclidian distances (Δϕ) between all modulatory states. Significance of differences between different modulatory states was determined with multivariate permutation tests.

To combine all dimensions, we plotted the one-dimensional projections of the Euclidian distances in the five-dimensional ϕ space (Δϕ) between all pairs of modulatory states (Fig. 4C), along with the results of statistical evaluation from multivariate permutation tests (see Methods). Decentralization significantly changed ϕ values from the intact state. In all three neuropeptides, ϕ values were different from the decentralized state, but not different from each other. Also, none of the neuropeptide states differed from the intact state, meaning that all of them essentially restored the intact values of ϕ.

The lack of differences in the pyloric phase relationships among different peptide bath applications goes contrary to the idea that neuropeptides with convergent cellular actions that target different subsets of circuit neurons give rise to different phase relationships (Marder and Weimann, 1992; Swensen and Marder, 2001). We addressed two concerns about this finding. First, we wanted to ensure that the lack of statistical difference did not arise from insufficient statistical power. We therefore included a larger data set from only extracellular recordings. Such recordings allow unambiguous determination of *F*_cycle_ and ϕ, but it is more difficult to obtain accurate spike counts (see Methods). Second, we wanted to ensure that consistent phase is not the automatic outcome of any neuromodulation that targets the pyloric circuit, independent of modulator identity and specific cellular and subcellular targets. We therefore included recordings obtained during bath application of DA (Fig. 5A). DA is not an unambiguously excitatory modulator, but rather has cell type-specific divergent actions on a range of ion channels and synapses (Harris-Warrick and Johnson, 2010).

**Figure 5:**
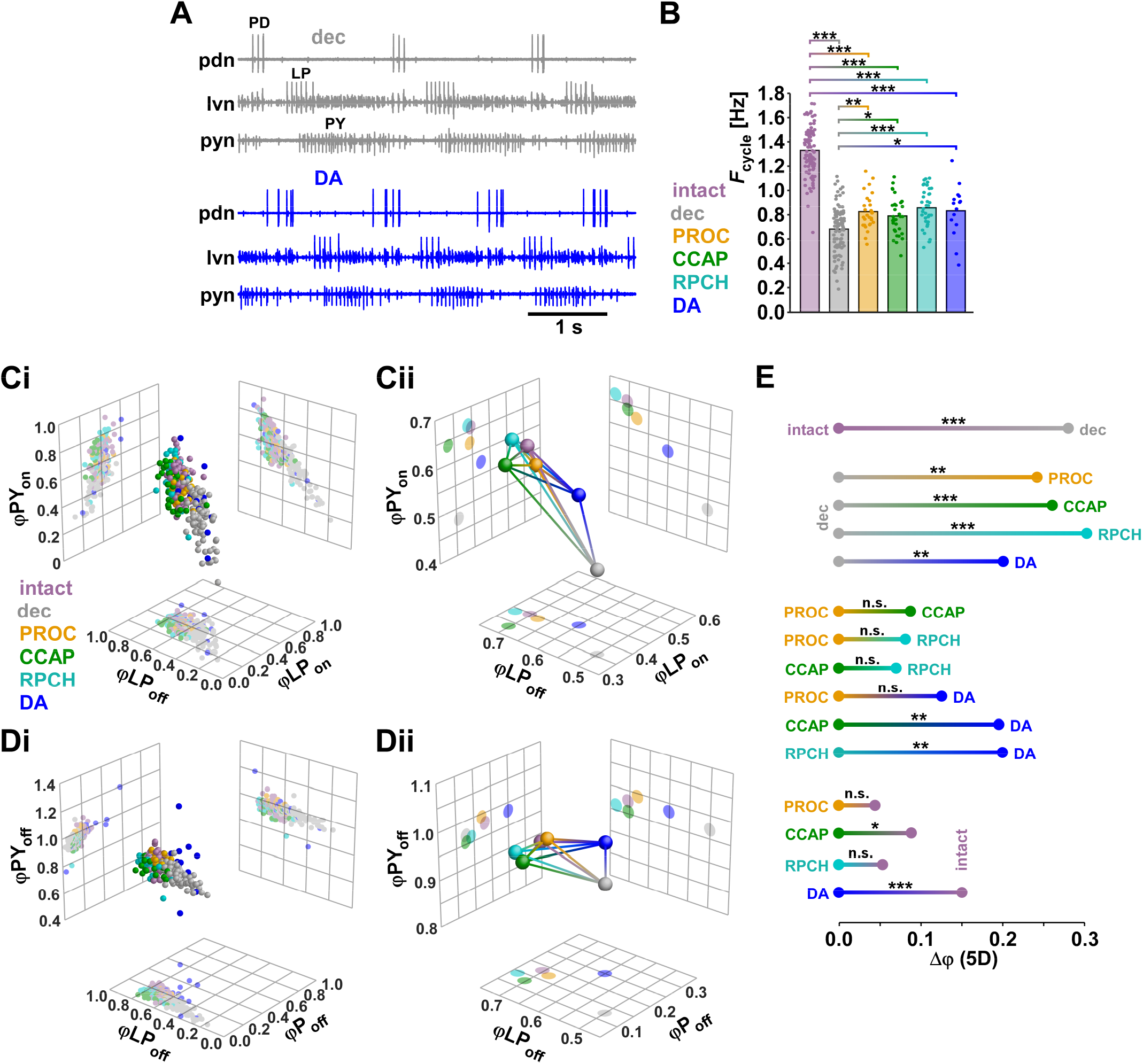
Analysis of *F*_cycle_ and ϕ in a large data set from extracellular recordings, including DA (intact: n = 105; decentralized: n = 100; PROC: n = 28; CCAP: n = 34; RPCH: n = 36; DA: n = 15). **A:** Example extracellular recording traces from a single preparation after decentralization and after application of DA. **B:** Means and individual data points for *F*_cycle_. One-way ANOVA showed signifcant difference between treatments (F_(5,317)_ = 150.7, p < 0.001, power = 1, normality and equal variance tests passed). Asterisks indicate significant difference determined by subsequent Holm-Sidak post-hoc tests. **C:** Three-dimensional plots (spheres) and corresponding two-dimensional projections (grids) of three of the five phases (ϕLP_on_, ϕLP_off_, and ϕPY_on_), for individual experiments (i) and centroids (ii). Lines connecting the centroids represent the Euclidian distances in three dimensions. **D:** Plots of a different combinations of three of the five phases (ϕPD_off_, ϕLP_off_, ϕPY_off_). **E:** One-dimensional projections of the five-dimensional Euclidian distances (Δϕ) between all modulatory states. Significance of differences between different modulatory states was determined with multivariate permutation tests.

In the larger data set, *F*_cycle_ increased from the decentralized state in all modulators, including DA (Fig. 5B). However, mean values were not substantially different from the smaller data set, and none of the modulators restored *F*_cycle_ to the level of the intact state. Three-dimensional plots of phases for individual data points (Fig. 5Ci, Di) and centroids (Fig. 5Cii, Dii) showed that intact and neuropeptide states cluster, while the DA state was more separated. DA responses also showed more substantial interindividual variability. Five-dimensional Euclidian distances (Δϕ) between all pairs of modulatory states were statistically evaluated (Fig. 5E). As expected, decentralization significantly changed ϕ values from the intact state, and ϕ values were different from the decentralized state in all modulators, including DA. Notably, consistent with the intracellular data shown in Figure 4, the neuropeptide states were not statistically distinguishable. However, DA was significantly different from CCAP and RPCH. While PROC and RPCH were not distinguishable from the intact state, the small difference between CCAP and intact was significant. Finally, the difference between DA and intact was highly significant. Therefore, we confirmed that phase relationships were not distinguishable between different neuropeptide states and that neuropeptides returned ϕ to values close to, or indistinguishable from, those in the intact state. These values were not the automatic outcome of modulation, as DA increased *F*_cycle_ from the decentralized state but elicited distinct phase relationships.

### Multidimensional comparison showed differences in spiking attributes between neuropeptides

Next, we analyzed the spiking behavior within bursts for the three neurons across all modulatory states, both for *#spks* (Fig. 6A) and *F*_spk_ (Fig. 6B). For visualization, we again plotted individual data points (Fig. 6Ai, Bi) and centroids (Fig. 6Aii, Bii). Compared to our findings for ϕ, in both sets of spiking attributes, modulated states appeared to cluster less, and the decentralized state was less separated from modulated states.

**Figure 6:**
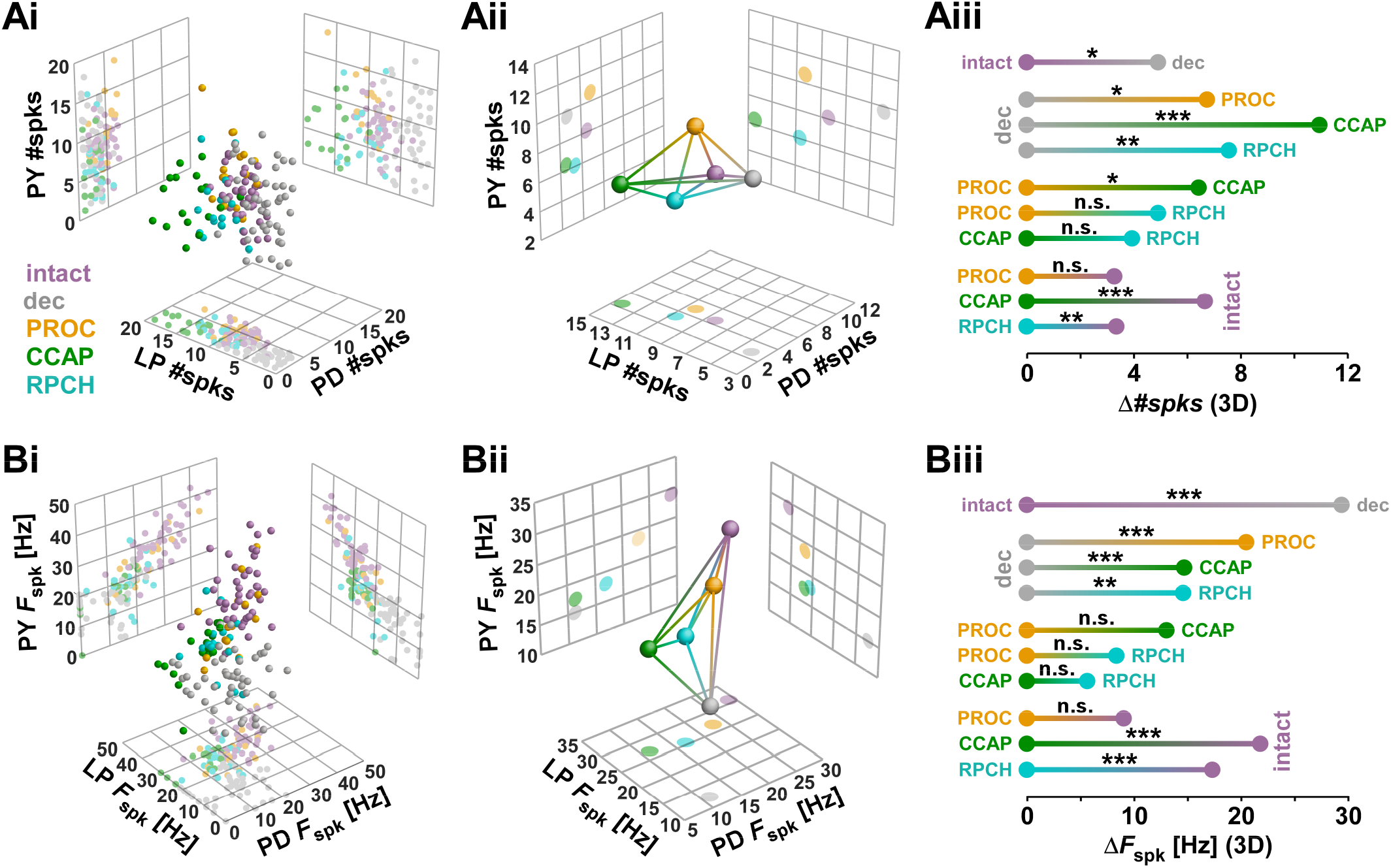
Euclidian distance analysis of spiking behavior. Data set is the same as shown in Figure 2. **A:** Analysis of number of spikes per burst in all three neurons.Three-dimensional plots (spheres) and corresponding two-dimensional projections (grids) for individual experiments (i) and centroids (ii). Lines connecting the centroids represent the Euclidian distances in three dimensions. One-dimensional projections of the Euclidian distances (Δ*#spks*) between all modulatory states (iii) are shown with the results of multivariate permutation tests. **B:** Same analysis as in **A**, but for spike frequencies within bursts in all three neurons.

Figure 6Aiii shows the one-dimensional projections of Euclidian distances in the three-dimensional attribute space for *#spks* (Δ*#spks*), along with the results of statistical evaluation. Decentralization significantly altered values of *#spks*, and application of all three peptides resulted in values different from the decentralized state. Across neuropeptides, PROC differed from CCAP, but RPCH was not statistically distinguishable from either PROC or CCAP. Both CCAP and RPCH were significantly different from the intact state, while PROC was not.

Figure 6Biii shows the one-dimensional projections of Euclidian distances in the three-dimensional attribute space for *F*_spk_ (Δ *F*_spk_), along with the results of statistical evaluation. As for *#spks,* decentralization significantly altered values of *F*_spk_, and application of all three peptides resulted in values different from the decentralized state. Across neuropeptides, no significant differences were found in the direct one-to-one comparisons. However, as for *#spks*, both CCAP and RPCH were significantly different from the intact state, while PROC was not. Therefore, overall, CCAP and RPCH elicited similar spiking behavior across the three different neuron types that was different from the intact state, while PROC essentially restored the spiking behavior of the intact state.

### Across all attributes, the output pattern in PROC can be distinguished from those in CCAP and RPCH

To include all attributes into a single analysis of differences across modulatory states, we rendered them unitless by *z*-scoring (Fig. 7A, see Methods). We then calculated the Euclidian distances in 12 dimensions (*F*_cycle_, 5 dimensions of ϕ, 3 dimensions of *#spks*, 3 dimensions of *F*_spk_). Figure 7B shows the one-dimensional projections of Euclidian distances in the 12-dimensional attribute space for *z*-scored values (Δ*z*-scores), along with the results of statistical evaluation. The decentralized state was significantly different from the intact state, and all neuropeptide states were significantly different from the decentralized state. Across neuropeptides, PROC was significantly different from CCAP and RPCH, but the difference between CCAP and RPCH was not significant. None of the neuropeptides fully recovered the output pattern, as all were significantly different from the intact state. This included PROC, which was indistinguishable from the intact state in the separate analyses of ϕ (Fig. 4C), *#spks* (Fig. 6Aiv), and *F*_spk_ (Fig. 6Biv). However, this is likely explained by the fact that the difference between PROC and the intact state was highly significant in the separate analysis of *F*_cycle_ (Fig. 2B).

**Figure 7:**
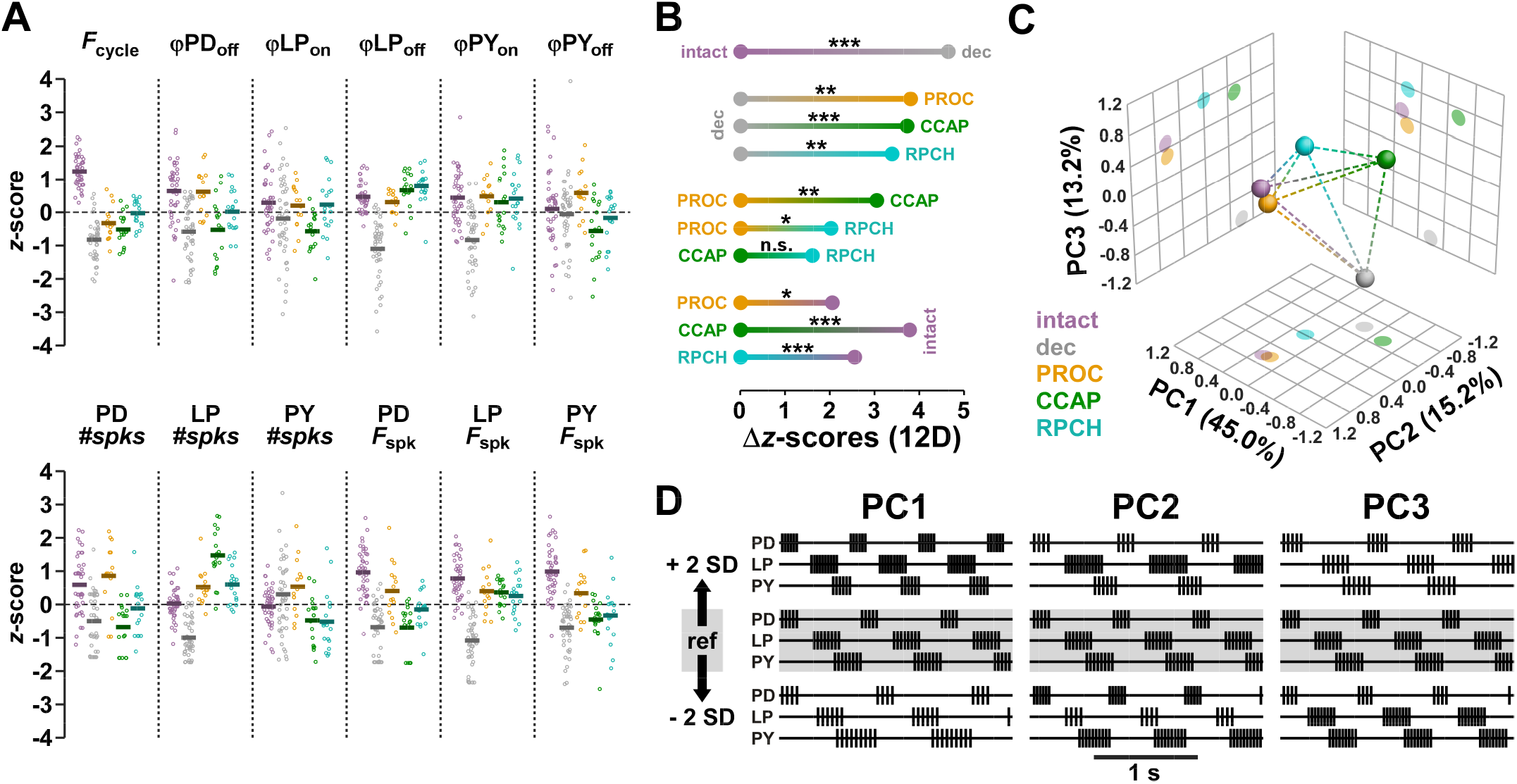
Euclidian distance analysis including all 12 output pattern attributes. **A:** Individual data points for all attributes, *z*-scored across all modulatory states. Means are shown as thick lines. **B:** One-dimensional projections of the 12-dimensional Euclidian distances (Δ*z*-scores) between all modulatory states. Significance was determined with multivariate permutation tests. **C:** Different modulatory states as centroids (spheres) and distances (dashed lines) in the dimensions of the first three principal components obtained by PCA. **D:** Theoretical change of output patterns along the first three principal components, in the range of ±2 standard deviations.

As an alternative way to visualize the differences between modulatory states, we performed PCA on the *z*-scored 12-dimensional data. In the dimensions of the first three principal components (Fig. 7C), which accounted for 73.4% of the variance, decentralization moved the output pattern to substantially different coordinates. Upon application of a modulator, PROC moved the pattern relatively close to the original intact state, while CCAP and RPCH were more distant from the intact state. Figure 7D depicts how the output pattern changes along those different principal components. Along PC1, the largest contribution to pattern changes was *F*_cycle_, followed by *F*_spk_ in all neurons and a subset of phases (ϕLP_off_, ϕPY_on,_ ϕPD_off_). Along PC2 and PC3, the largest contributions were from LP *#spks* and ϕLP_on_, respectively.

From all separate and joint analyses of output pattern attributes, we conclude that CCAP and RPCH elicit similar output patterns in the pyloric circuit, while PROC elicits a distinct pattern that is closer to the intact state than the ones elicited by CCAP or RPCH.

### Identification of output patterns by machine learning algorithms was only moderately successful

All analyses shown so far were based on attributes that are well-established for the quantification of rhythmic circuit output but are ultimately an arbitrary subset of measures that can be taken from the recording traces. We therefore asked the question how well patterns could be distinguished with a less biased approach. To this end, we employed machine learning algorithms that were trained on a subset of data from each modulatory state and then used to blindly classify output patterns by modulatory state. The general idea was to ask how successful the algorithm is in correctly identifying a modulatory state across a selection of given individual output patterns.

For a direct initial comparison with our attribute-based approach, we first used a pattern recognition network (Fig. 8A) that solely relied on the same selected attributes we extracted from the recordings, namely the 12-dimensional *z*-score values used in Figure 7. Figure 8B shows the classification matrix from this approach. A high percentage of data from the intact and decentralized states were correctly classified, with error rates of <10%. The algorithm also classified most neuropeptide states correctly but had a much higher error rate for each. Misclassifications corresponded reasonably well with our results from determining Euclidian distances in some cases, but less so in others. CCAP and RPCH (which were statistically indistinguishable) had largest percentages of misclassifications as each other, meaning these two states were easily confused. PROC was often misclassified as the intact state (which it was statistically indistinguishable from), but also as RPCH.

**Figure 8:**
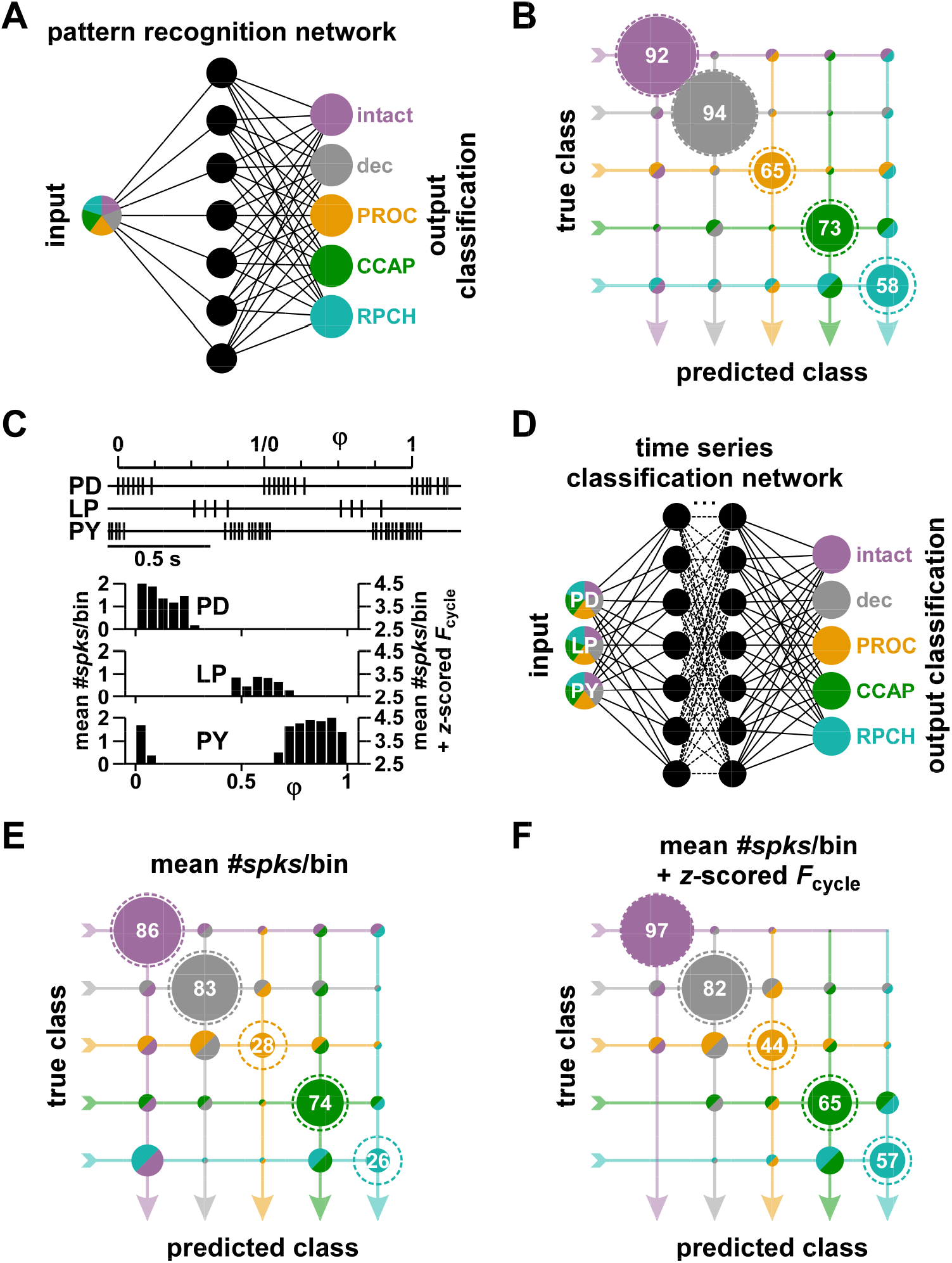
Classification of modulatory states with machine learning algorithms. **A:** Schematic of the pattern recognition network architecture (see Methods) trained to classify modulatory states from the same circuit output pattern attributes used for the 12-dimensional Euclidian distance analysis shown in Figure 7. **B:** Classification matrix of the results from the pattern recognition network. The true class is the actual modulatory state and the predicted class the identification result. Percentages of predicted classes for each true class are represented by the areas within circles. The number values of correct classifications represent the percentages along each true class row. Areas inside the dashed circles represent 100%, the size of the circle if all experiments were classified correctly. Because of the unequal numbers of experiments in each true class, the circle areas add up to 100% only within each row. **C:** Example of spike timing of PD, LP, and PY in two consecutive normalized cycles, and example spike phase histogram from mean spike counts across all cycles in a 30s recording. Mean *#spks*/bin were both used as they were (left scale) and after adding the *F*_cycle_ *z*-score value (right scale). **D:** Simplified schematic of the time series classification network architecture (see Methods) trained to classify modulatory states from the spike phase histograms. **E:** Classification matrix from phase histograms of mean *#spks*/bin values. **F:** Classification matrix from phase histograms of mean *#spks*/bin values offset by *F*_cycle_ *z*-score values.

To rely less on predetermined attributes, we performed a more minimal analysis by extracting spike times of each neuron and sorting them into a fixed number of bins per cycle. Figure 8C shows an example of a spike phase histogram constructed with this approach. Because solely relying on *#spks*/bin does not account for differences in *F*_cycle_ between modulatory states, we subsequently added the *z*-scored *F*_cycle_ values to the bin counts. We used these data to train a time series classification network (Fig. 8D). Overall, classifications were less successful than with the pattern recognition network. Furthermore, error rates were larger when we used only the *#spks*/bin values (Fig. 8E) than when we added *z*-scored *F*_cycle_ values (Fig. 8F). With *F*_cycle_ included, classifications were highly successful for the intact state, and still had a relatively low error rate for the decentralized state. PROC had the highest error rate and curiously was more often confused with the decentralized state than the intact state. Again, the largest percentage of misclassifications for CCAP and RPCH was as each other, meaning these two states were still easily confused. Except for PROC, the algorithm recognized the correct modulatory state in the majority of cases, but also had relatively high error rates for the other two neuropeptides. Overall, the findings of the two ML analyses are consistent with our Euclidian distance analysis.

## Discussion

### Cellular convergence of modulator actions as the predominant mechanisms of circuit activation

Different neuromodulators converging on the same intracellular targets can have distinct actions at the circuit level if the circuit neuron types have different receptor expression patterns for these modulators. Swensen and Marder (2001) showed such circuit-level divergence by mapping the neuropeptide activation of *I*_MI_ across pyloric neuron types (Fig. 2A). Our finding that circuit activity in PROC is most similar to the intact state, and that activities in CCAP and RPCH are statistically indistinguishable, match well with these circuit activation maps. However, overall, the differences between modulated states were more subtle than expected, particularly because the activation maps are a simplified representation of neuropeptide actions. While neuropeptides can converge to activate *I*_MI_ (Swensen and Marder, 2000, 2001), the magnitude of activation can differ (Swensen and Marder, 2000; Garcia et al., 2015; Li et al., 2018), and *I*_MI_ is now known to consist of at least two components that could be differentially activated (Rodriguez et al., 2013; Schneider et al., 2021). In addition, neuropeptides influence synaptic efficacy and dynamics in the STG (Thirumalai et al., 2006; Atamturktur and Nadim, 2011; Zhao et al., 2011; Garcia et al., 2015), and synaptic effects may be distinct (Li et al., 2018).

We found that differences between modulated states were mostly found in spiking activity but not burst phases. With intact descending inputs, burst phases remain relatively independent of *F*_cycle_ across animals (Fig. 3A) (Bucher et al., 2005; Goaillard et al., 2009; Anwar et al., 2022). As reported previously, (Luther et al., 2003), we showed that phase constancy requires the presence of neuromodulators, as decentralization changed the phase relationships and eliminated phase maintenance across *F*_cycle_ values. All neuropeptides recovered the phase relationships and phase constancy seen in the intact circuit. This was particularly surprising because the intact circuit is under control of many modulators, including non-peptides (Marder and Bucher, 2007), and effects of different modulators can differ quantitatively and are not necessarily additive (Garcia et al., 2015; Li et al., 2018). A possible reason is that pyloric phases are mainly determined by the pacemaker neurons. LP and PY burst on rebound from pacemaker inhibition, and it has been suggested that the voltage-dependence of *I*_MI_ results in rebound timing that changes proportionally to the duration of inhibition (Schneider et al., 2021). As all three peptides act on the pacemaker neurons and produce similar values of *F*_cycle_ and ϕPD_off_, there would be little effect on rebound phase. This does not mean that the activity phases of follower neurons are altogether unaffected by neuromodulation, as monoamines have a strong effect on these phases (Harris-Warrick and Johnson, 2010), which we confirmed with DA application (Fig.5).

In contrast to burst phases, spiking activity was substantially affected. All three peptides target AB and LP, but only PROC in addition targets PD and PY, which is reflected in higher values of PD and PY *F*_spk_ in PROC (Fig. 2E). This is consistent with results from Swensen and Marder (2001), who demonstrated that the dynamic clamp addition of *I*_MI_ to PD and PY in CCAP was sufficient to increase their spiking to the levels seen in PROC. The fact that these neuropeptides neither restored the spiking behavior nor *F*_cycle_ seen in the intact state (Fig. 2B, 5B) may be due to the presence of additional modulators when descending inputs are intact.

### Multi-dimensionality of circuit outputs

We present a novel way of determining differences in circuit outputs, collapsing information from multiple output attributes into a single measure. Because output attributes have different units, we used *z*-scored values to perform higher-dimensional analyses. The multivariate permutation test provided a reliable statistical comparison between the *z*-scored vectors (Anderson, 2001; Nichols and Holmes, 2002), but it does not explicitly quantify the magnitude of differences. To measure differences, we used the Euclidean distances of the population centroids, which provides a geometric description in cases where circuit activity changes across different conditions. Other methods such as nonlinear unsupervised dimensionality reduction (Gorur-Shandilya et al., 2022) may constrain the differences to fewer dimensions but would not provide an explicit measure of difference.

Our approach has two caveats. First, all attributes are equally weighted. This does not mean that all attributes are equally important for motor activity, but we argue that an unbiased analysis is warranted when there is no *a priori* way of knowing which attributes are the most salient. For example, Daur et al. (2021) showed that in one muscle innervated by PD, responses to bursting input were moderately sensitive to changes in *F*_cycle_ or PD *F*_spk_, while another muscle was insensitive to *F*_cycle_ but acutely sensitive to PD *F*_spk_. Second, the information about the specific attributes that contribute to the differences is lost. Therefore, our method does not eliminate the need to also analyze attributes individually or perform principal component analysis.

We used a subset of commonly quantified output attributes. In a recurrent circuit, all output pattern attributes are potentially interdependent, and one must therefore gauge the degree of redundancy. For example, we did not include latencies, which were highly correlated with *P*, but included phases which were not (Fig. 3A). Ultimately, any selection of attributes is subjective and additional attributes could reveal differences in output patterns. For example, motor patterns can show substantial cycle-to-cycle variability (Horn et al., 2004; Bucher et al., 2005), which can be important for circuit function and behavior (Brezina et al., 2006). In the pyloric circuit, cycle-to-cycle variability depends on inhibitory synaptic feedback to the pacemaker neurons (Nadim et al., 2011) and decreases when that feedback is enhanced by neuropeptides like PROC (Zhao et al., 2011). Neuromodulators can also change the precise interval structure of spikes within each burst instead of just the mean *F*_spk_, with potential consequences for rhythm generation and muscle activation (Szucs et al., 2005; Ballo et al., 2012; Daur et al., 2021). Furthermore, we only used the core pyloric circuit for our analyses. In their study of *I*_MI_ activation by different neuropeptides, Swensen and Marder (2001) included the inferior cardiac (IC) and ventral dilator (VD) neurons, which participate in both pyloric and gastric mill circuit activity. PROC activates *I*_MI_ in both VD and IC, CCAP activates *I*_MI_ in IC, while RPCH does not target either. The STG also includes the neurons of the gastric mill circuit which interacts with the pyloric circuit (Weimann et al., 1991; Bucher et al., 2006; Marder and Bucher, 2007; Daur et al., 2016), especially through actions of neuromodulators (Wood et al., 2004). It is possible that inclusion of a larger set of circuit neurons and additional activity attributes would reveal more differences between circuit outputs, but such differences would likely still be small compared to differences between neuropeptide- and monoamine-elicited patterns (Marder and Weimann, 1992).

### Do convergence and similar circuit outputs suggest redundancy?

The observation that different neuropeptides elicit similar circuit outputs through convergent cellular actions suggests redundancy in the neuromodulator complement. This may not be surprising, as there are more than 100 neuropeptides found in a crustacean species, although many of them are peptide family isoforms that can have shared actions (Dickinson et al., 2016). However, the similar output patterns we describe here were elicited in response to bath applications and do not encompass more nuanced actions of modulators released by projection neurons, which can be differentially activated across different contexts and with different cotransmittters (Nusbaum et al., 2001). In addition, differences between output patterns may be unmasked by responses to additional inputs. For example, two very similar gastric mill circuit activity patterns become distinct in response to additional hormonal or sensory inputs (Powell et al., 2021). Therefore, even the statistically indistinguishable patterns in CCAP and RPCH may respond differently to the same physiological perturbation.

Furthermore, circuits are likely under the influence of multiple comodulators at any time (Nusbaum et al., 2001; Nadim and Bucher, 2014), and it matters how they interact. Two neuropeptides with convergent actions may have only partially occluding effects because of differences in the expression levels of their receptors and nonlinear substrate interactions. For instance, saturating CCAP concentrations only occlude the effect of PROC in LP but not IC, even though both modulators target both neurons (Garcia et al., 2015). Additionally, the combined effects of PROC and CCAP on *I*_MI_ become nonlinear if either modulator is applied below saturation levels (Li et al., 2018). Therefore, even if two modulators with convergent cellular actions act on an identical subset of circuit neurons, comodulation with both may give rise to circuit activity different than that elicited by each individual modulator.

### Interindividual variability

Our finding of similar output patterns must be viewed in the context of population variability. Our statistical comparisons were done with unpaired experimental paradigms, considering each modulatory state as a separate population. It is possible that output patterns across different neuropeptides would be better distinguishable in individual preparations and that interindividual variability masked differences in our analysis. For example, Swensen and Marder (2001) performed paired comparisons from sequential applications of PROC and CCAP in each experiment and found significant differences in ϕPD_off_ and ϕPY_off_ that we did not detect here. Paired comparisons have greater statistical power in testing whether changing the modulatory state has consistent effects across individuals. However, we were also interested in a conceptually related but different question, i.e., whether a specific modulator elicits characteristic activity that can be identified across individuals. For this question, the unpaired paradigm is arguably more appropriate, and we found that identification of characteristic patterns was unreliable.

What is the source of variability in circuit outputs? Ionic currents can vary substantially across individuals, but compensatory tuning rules are thought to produce unique solutions to generating similar output activity in individuals (Marder and Goaillard, 2006; Calabrese et al., 2011; Turrigiano, 2011; Golowasch, 2019a; Goaillard and Marder, 2021). Neuromodulation can reveal differences in parameter sets but can also be integral to the tuning rules that produce similar outputs and therefore counteract variability (Hamood and Marder, 2014; Marder et al., 2014a; Marder et al., 2014b; Golowasch, 2019b; Goaillard and Marder, 2021; Maloney, 2021; Tamvacakis et al., 2022). In the STG, neuropeptide receptor expression can vary substantially (Garcia et al., 2015), and both receptor and ion channel expression levels depend on neuromodulator presence (Temporal et al., 2012; Lett et al., 2017). Neuropeptide modulation can reduce interindividual variability in single neuron response properties (Schneider et al., 2022) and of circuit outputs (Schneider et al., 2023). The distribution of *z*-scored attribute values in our data also suggests that variability in at least a subset of attributes is reduced compared to the decentralized state (Fig. 7A), while it remains substantial enough to prevent unambiguous output classification.

Although we focused on a small invertebrate pattern-generating neural circuit, both cellular-level variability of ion channel and receptor expression (see above) and interindividual variability of network activity and behavior are common across neural circuits and animals (Feierstein et al., 2015; Xu et al., 2021; Cerins et al., 2022; Rihani and Sachse, 2022; Tamvacakis et al., 2022; Moujaes et al., 2023). Understanding neuromodulator effects at the population level also has important health ramifications (Shamay-Tsoory and Abu-Akel, 2016; Pavăl and Micluția, 2021; Moncrieff et al., 2022), because almost all therapeutic drugs used to treat mental disease and disorders influence natural modulatory processes. Therefore, our findings have broad significance for understanding neuromodulatory actions in face of interindividual variability both for understanding normal physiological function and in understanding drug actions in the brain.

## Acknowledgements

We thank Gareth Russell for advice on statistics, and Omar Itani, Jorge Golowasch, and Dawn Blitz for helpful suggestions and comments.

Current affiliation of ACS: FB10 Animal Physiology, University of Kassel, Kassel, Germany.

## Extended Data Legends

**Figure 3-1:**
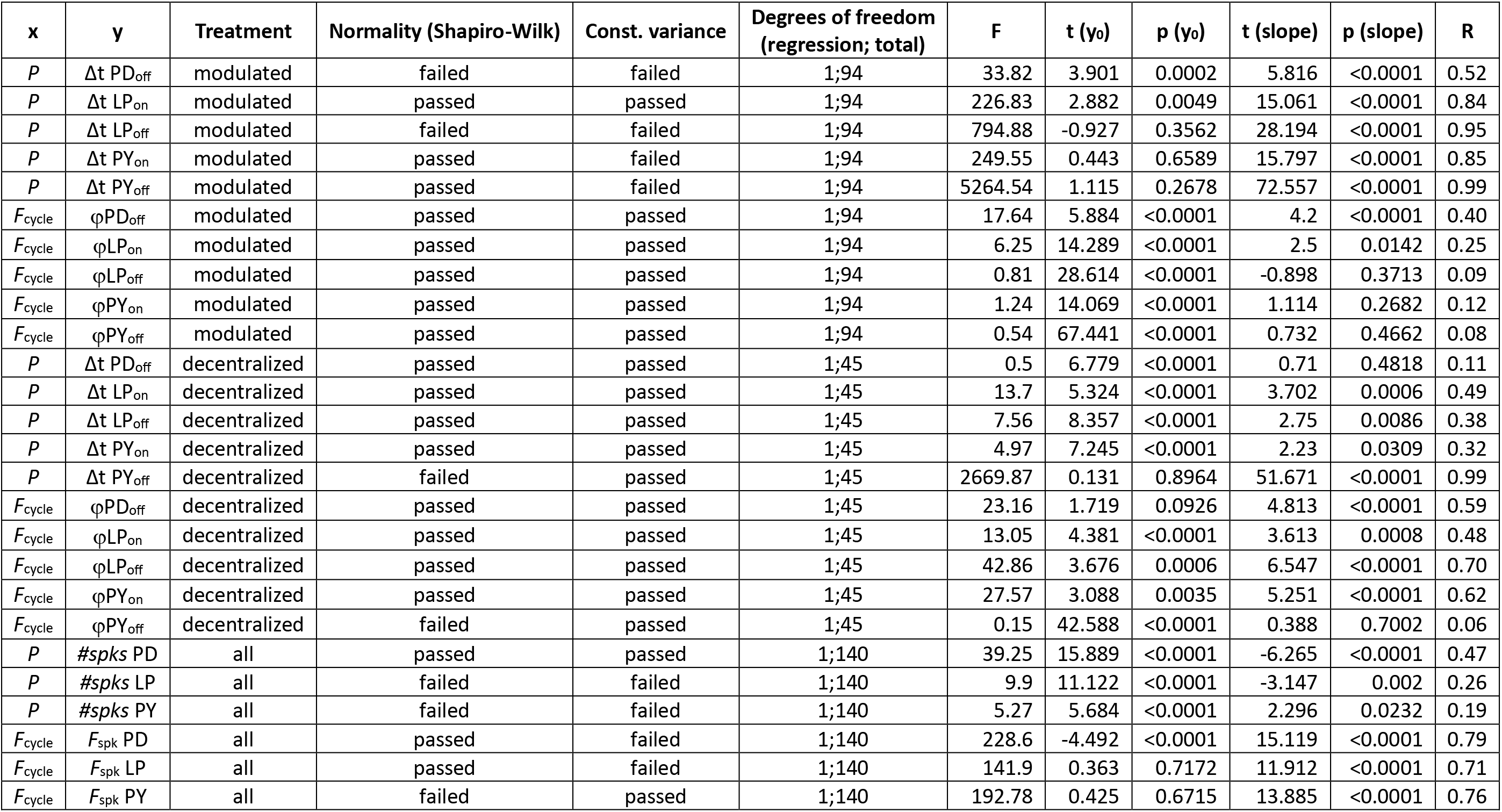
Regression parameters for the analyses shown in Figure 3.

